# Spatiotemporal changes in soil properties predict plant health at continental scale

**DOI:** 10.1101/2025.01.07.631818

**Authors:** Ningqi Wang, Gaofei Jiang, Yugang Hou, Changqin Li, Yuanduo Meng, Peijie Chen, Yuling Zhang, Tianjie Yang, Xiaofang Wang, Xinlan Mei, Thomas Pommier, Samiran Banerjee, Matthias C. Rillig, Francisco Dini-Andreote, Ville-Petri Friman, Yangchun Xu, Alexandre Jousset, Qirong Shen, Zhong Wei

## Abstract

Abiotic and biotic soil properties are strong predictors of plant yield globally^1–5^, but they become unreliable over large areas when plant health is threatened by pathogen^6–8^. Here we present a novel approach to predict plant health based on spatiotemporal changes in soil chemical and biological properties. We first demonstrate that plant health and soil properties consistently respond to environmental change (organic fertilization) regardless of the soil type or geographical origin. Second, we experimentally show that trackable shifts in soil properties reliably explain soil suppressiveness to the *Ralstonia solanacearum* bacterial pathogen and that a scalable spatiotemporal model predicts plant health with 84% accuracy across multiple climatic zones and cropping systems. Our results suggest that this tight coupling between soil properties and plant health could facilitate the development of agricultural practices aimed at sustainably improving crop yields while safeguarding crop health.

## Main

Soil physicochemical and microbial properties are strong predictors of plant yield globally^1–3^, but they become unreliable under disease pressure. Predicting crop vulnerability to pathogens and pests remains difficult because we poorly understand which soil properties determine plant health^4,5^ (Fig. 1a). As plant diseases and pests cause around 30% of annual crop losses globally (FAOSTAT, Data of Access: [10-12-22]), harnessing the potential of soils to prevent disease is essential for designing low-input and high sustainable agroecosystems toward sustainable intensification goals ^6,7^. Soil chemical and biological properties supporting plant health can be modulated by informed agricultural practices, such as the use of organic fertilizers, which have a long history of improving plant health sustainably ^8^. However, organic fertilizers have only resulted in enhanced plant health in less than one-third of the cases^9^, highlighting the difficulty in consistently producing the expected benefits for crop protection.

**Fig. 1:**
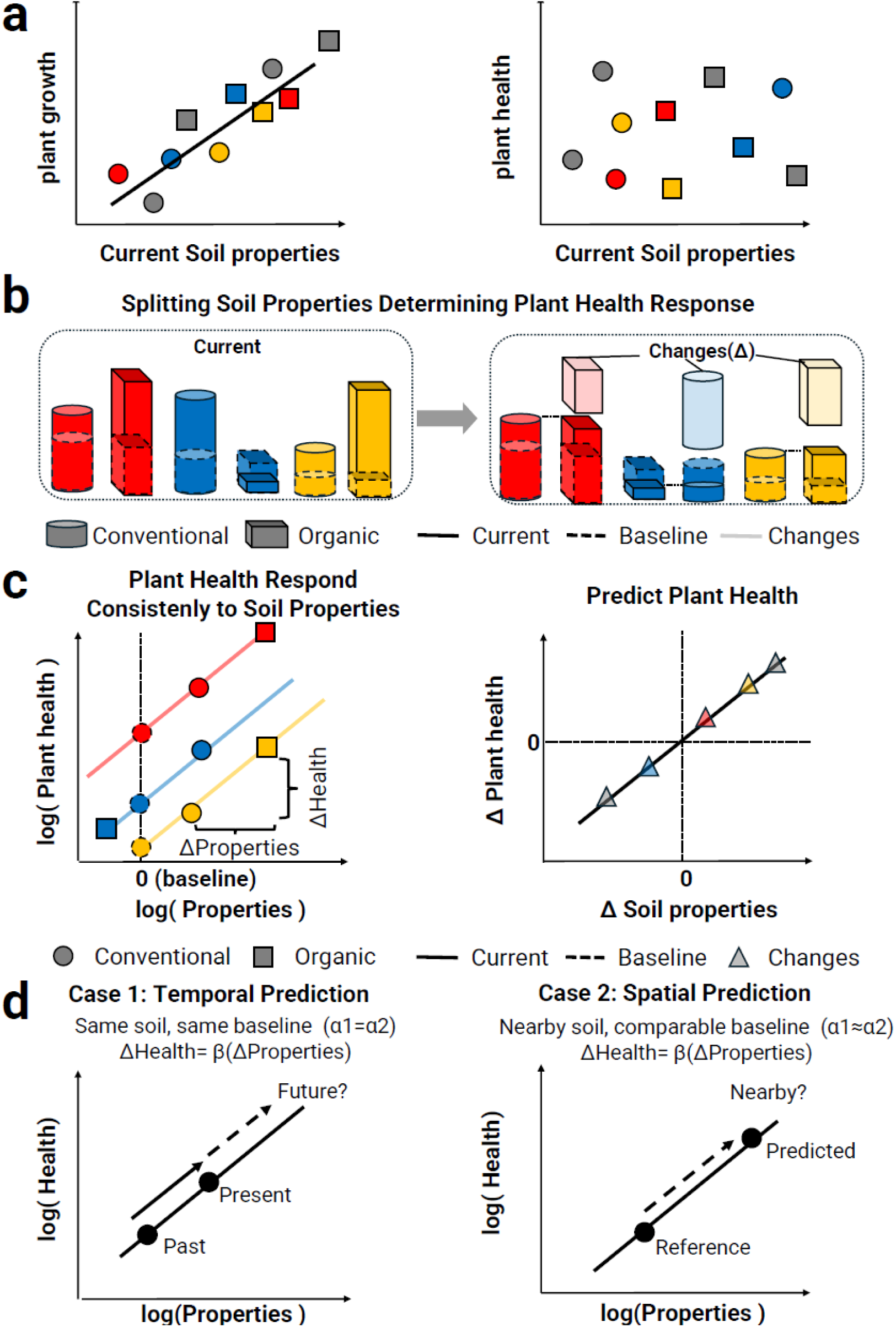
Conceptual overview of the spatiotemporal model for global plant health prediction. **a: Challenges for predicting plant growth and health across large spatiotemporal scales.** Plant growth can be predicted based on soil properties because soil fertility directly determines soil nutrients available for supporting plant growth. By contrast, using soil properties to predict plant health remains a challenge because plant health is determined by multiple interacting factors that vary spatially and temporally. **b, Partitioning of soil properties into baselines and changes modulating plant health.** Each type of soil has a unique baseline of soil properties hidden within the snapshot measurement, which disrupts predictions of plant health. To eliminate the impact of varied baselines, we split changes in soil properties determining plant health response by comparing pairwise soils with the same baseline. The log-scale proportions between pairwise soils were calculated and defined as changes (Δ) modulating plant health. **c, Changes of soil properties as a predictive toll of plant health.** Assuming changes in plant health respond consistently to changes in soil properties. Thus, each soil exhibits a parallel response in trajectories in soil property plant health space (log scale). Once the response of plant health to changes in soil properties is consistent across sites, plant health can be expressed as a linear function of soil properties changes, thus enabling prediction. d, Use cases of plant health prediction. Case 1: For the soils with the same histories (i.e. same baseline), future plant health development can be inferred according to past dynamics in soil properties. Case 2: For nearby soils, the baselines are similar, allowing for direct comparison in soil properties for plant health prediction. Note that both soil properties and plant health are expressed on a log scale and that the regression coefficients are derived from the log ratio. Shapes denote different management practices, and colors denote different sites, dash lines denote baseline. Specifically, the triangle denotes changes between different management practice.

Soil suppressiveness, defined as the natural ability of some soils to prevent diseases ^10^, is one of the most important aspects of plant health. The challenges in explaining and determining soil suppressiveness are two-fold. First, plant disease outbreaks are caused by multiple interacting factors and identifying universal key abiotic or biotic drivers has proven challenging ^8,11^. Second, the relative importance and interactions between soil properties vary across small geographical and temporal scales, making it difficult to extrapolate observational patterns observed at one location across broader spatial and temporal scales ^12,13^. As a result, while single time-point measurements (henceforth *“snapshot”*) of soil chemical or biological properties may explain soil suppressiveness locally ^14,15^, they often fail to do so at broader geographic and temporal scales mostly due to increased environmental heterogeneity and variability ^16,17^.

Here we present a new analytical approach reliably predicting plant health based on measured soil properties across multiple samples from different sites. This approach draws inspiration from previous studies demonstrating that system-level processes spanning from human health ^18^ to ecosystem resilience ^19,20^, are best predicted by the rate of change rather than by single snapshot measurement of system properties ^21^. We first separated site-specific baseline and changes in soil properties induced by management (here organic versus conventional farming, figure 1b). Consequently, while baseline soil properties vary between locations, plant capacity to respond to changes in soil properties may be conserved, allowing us to predict plant health as a functional response to environmental change. Based on the assumption that the responses to changes in soil properties are consistent across sites, plant health is expected to change along the log scale despite differences in site-specific baselines. This would facilitate expressing plant health as a linear function of changes in soil properties (Fig. 1c) to predict future plant health trajectories, infer the suppressiveness of unknown soils, and estimate the costs and benefits of soil restoration efforts (Fig. 1d).

To predict plant biomass and health based on soil properties at a large geographic scale, we collected 181 agricultural soil samples from fields across tropical to subarctic regions in China (Extended Data Fig. 1a and Supplemental Table 2). We measured ten key soil chemical and microbial properties and evaluated tomato biomass (dry weight) and plant health (pathogen density and disease severity) when challenged with the pathogen *Ralstonia solanacearum*. This pathogen ranks among the top ten of the most devastating plant pathogens capable of infecting over 250 plant species ^22^ (Supplemental Table 2). While three out of ten soil properties, including soil total nitrogen content, could adequately explain plant biomass (explanatory power =31%), these parameters were insufficient to predict changes in pathogen densities and disease severity (explanatory power <5%, Extended Data Fig. 1-2 and Table 3), demonstrating the challenge of predicting plant health based on soil properties collected at a single time point.

To test our analytical approach for predicting plant health, more specifically soil disease suppressiveness, we first conducted a global meta-analysis to determine whether soil responses to organic and conventional farming practices are spatially conserved. Using locally paired soil samples exposed to organic or conventional fertilization management, we explored whether the observed patterns in plant health can be inferred from changes in soil properties (Fig.2a and Extended Data Fig.3 and Table 4,5). Organic fertilization not only shifted multiple soil chemical and biological properties, particularly increasing bacterial diversity but also enhanced plant health by reducing disease severity and pathogen density (Fig. 2b). However, due to the incompleteness of variables included in the publicly available datasets, our analysis failed to link the plant health with changes in soil properties (Fig. 2a Venn diagram and Table 5). Nevertheless, we found that the response of soil microbiological and chemical parameters to organic fertilization was highly conserved despite large, site-specific changes in soil property baselines (Extended Data Fig. 4-5 and Table 6).

**Fig. 2:**
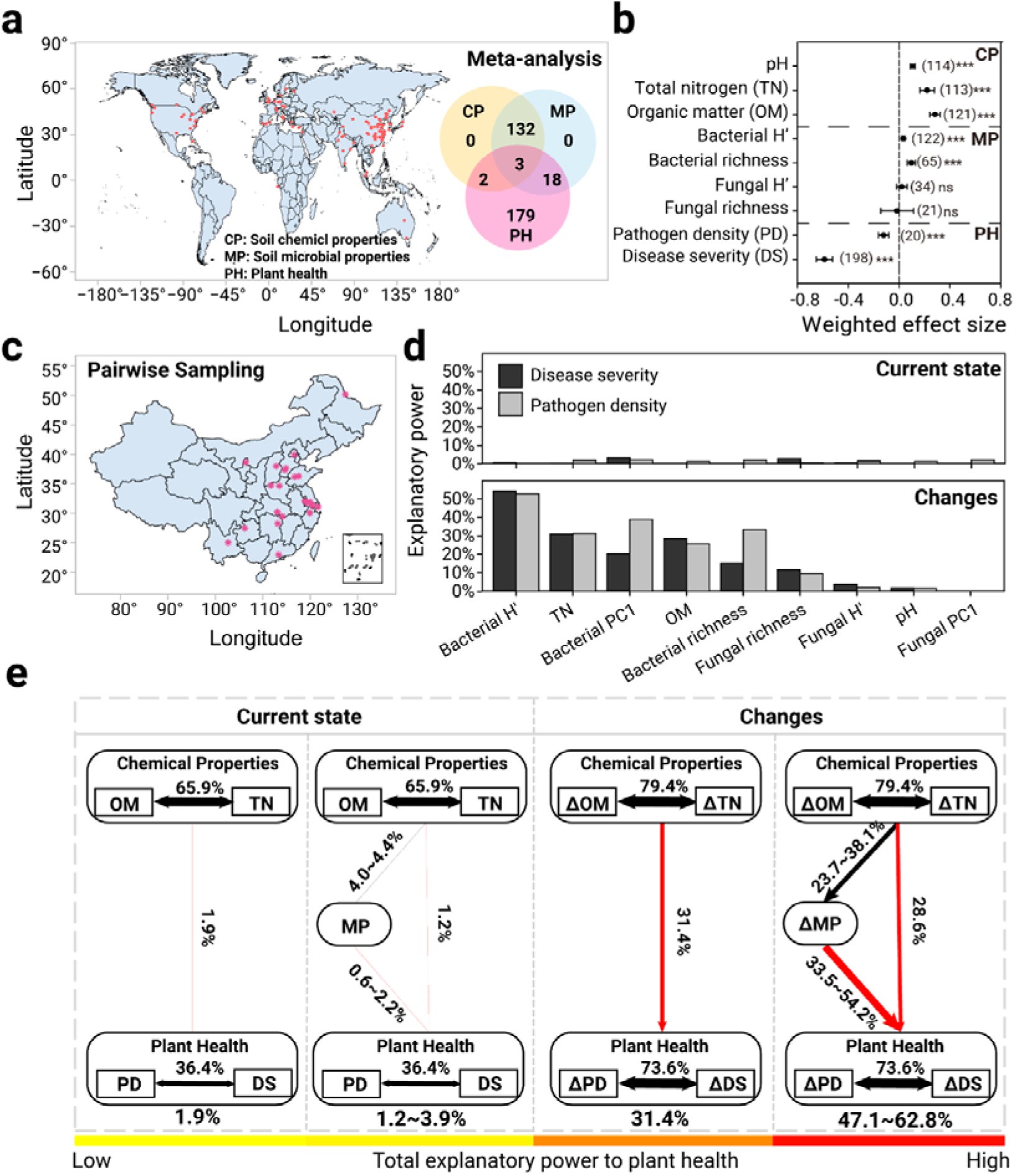
Spatiotemporal dynamics of properties between pairwise soils predict plant health. **a: Geographical overview of the covered studies investigating the effect of fertilization regime on soil properties and plant health.** Each point corresponds to one study containing pairs of samples with a recorded history of diverging organic and conventional fertilization regimes (≥3 years). The Venn diagram on the right highlights the core number of observations covering soil chemical properties (CP: pH, soil nitrogen content, and organic matter content defined as organic carbon content), microbial properties (MP: bacterial and fungal diversity), and plant health (PH: pathogen density or disease index). **b: Weighted effect size of organic fertilization regime on soil chemical properties, microbial properties, and plant health.** Positive values indicate a positive effect of organic fertilization on the tested parameter relative to conventional fertilization. Error bars represent 95% confidence intervals. Numbers in brackets indicate the number of experiments. Asterisks represent the significance levels. ns: non-significant, * *p* < 0.05, ** *p* < 0.01, *** *p* < 0.001**. c: Geographical overview of collected pairwise soil sampling connecting soil properties and plant health.** Each point corresponds to one site containing pairs of collected soil samples with a recorded history of diverging organic and conventional fertilization regimes (≥3 years). All collected soil samples were measured for soil chemical properties (pH, soil nitrogen content, and organic matter content), microbial properties using metagenomic analysis (bacterial and fungal diversity) and plant health (their resistance against *R. solanacearum* measured by pathogen density and disease index). **d: Explanatory power of soil chemical and microbial properties to plant health.** Current properties do not explain plant health, while changes within each pair robustly predict changes in plant health. **e: Structural equation modelling highlighting the key relationships between chemical properties, microbiome properties, and plant health**. The left part shows the effect regarding snapshot measurement (current state), and the right part shows the effect regarding changes. Percentages above each arrow and the arrow size represent the explanatory power of each path. The range of explanatory power was summarized from SEMs with different microbial properties included, such as bacterial richness, PC1, and the Shannon index (See detailed SEMs in Extended Data Fig. 8). Black and red arrows indicate positive and negative correlations, respectively. The bar at the bottom represents the total explanatory power of the SEM. Abbreviations, CP: soil chemical properties, MP: soil microbial properties, PH: plant health, TN: soil total nitrogen content, OM: soil organic matter content, H’: Shannon index, PC1: first axis of the PCA analysis on bacterial community, DS: disease severity, PD: pathogen density, Δ: changes, defined as the log-ratio proportion of a given parameter between organic and conventional fertilization.

To experimentally test whether the shifts in soil microbial diversity and chemical properties predict plant protection against the *R. solanacearum* (Fig. 2c-e), we collected pairwise soil samples from organically and conventionally fertilized agricultural fields across 23 agricultural sites in China (ranging from tropical to subarctic regions, Fig. 2c) with high-quality metadata on soil chemical and biological properties (Supplementary Table 7). We then used a large-scale and controlled greenhouse experiment to establish a direct link between plant health and changes in soil properties when exposed to infection by *R. solanacearum*. Organic fertilization showed a site-specific effect on soil chemical properties and plant health (Extended Data Fig. 6), decreasing both disease severity and pathogen densities (Extended Data Fig.7). While current soil properties explained only 2.0-3.3% of the plant health (best five predictors, Fig. 2d, “Current” panel, Supplementary Table 8), we could account for up to 54.2% of plant health (with Bacterial H’ having the best explanatory power, followed by soil total nitrogen at up to 30%) based on changes in soil properties changes. This confirms that plant health consistently responded to changes in soil properties, with microbial properties playing a critical role (“Changes” panel, Fig. 2d, Supplementary Table 8).

To disentangle the direct and indirect effects of soil chemical and biological properties on the plant health, we employed structural equation modelling (SEM). The explanatory power of the models increased from <2% to up to 30% when changes in soil properties were included (Fig 2e, left panel), and a further increase of up to 63% was achieved when changes in both soil chemical and microbial properties were integrated (Fig 3e, right panel, Supplemental Fig 8-9, and Table 9). Microbial properties had the highest direct effect on plant health (Extended Data Fig. 7 and 8), and nearly one-third (30.4%) of all the identified bacterial genera showed significant correlations with both pathogen density and disease severity (Supplementary Table 10), highlighting the importance of the microbiome for soil suppressiveness. We further performed a stepwise regression modelling to select the most parsimonious set of soil properties to predict plant health. Compared to the current soil properties (prediction accuracies of 22.3% for disease severity and 7.2% for pathogen density), changes in soil parameters increased the prediction accuracies of 72.1% for disease severity and 63.2% for pathogen densities (Extended Data Fig. 10). These results demonstrate that plant health can be accurately predicted based on soil responses to organic fertilization.

**Fig. 3:**
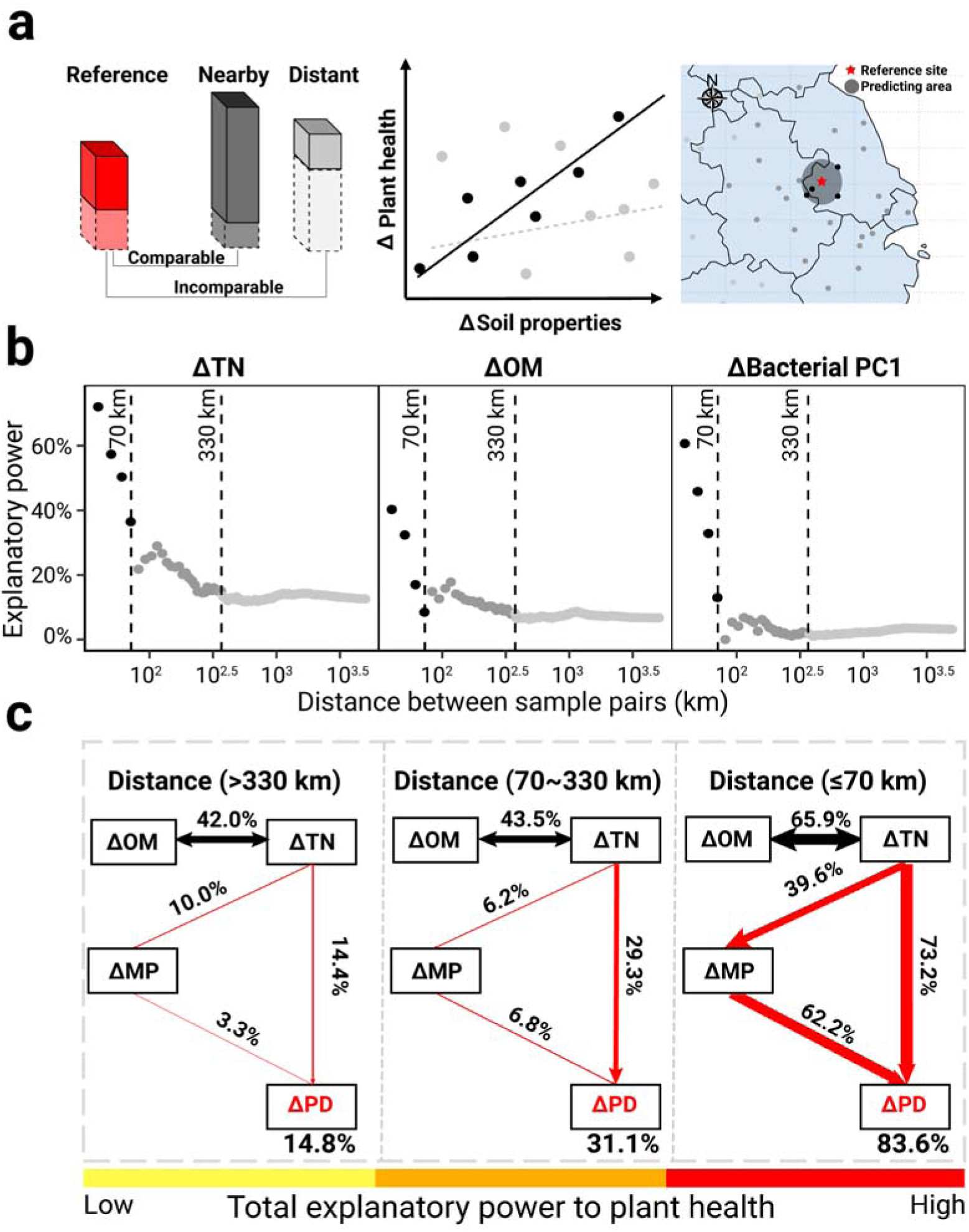
Spatial extrapolation of plant health prediction based on changes in soil chemical and microbial parameters from a broad sampling survey across various climates, soil types, crop history, and biomes. **a: Conceptual overview of the large scale spatial model for plant health prediction**. Each reference site has a surrounding area within which soil health can be predicted with high accuracy. We assume that within such areas (dark gray circles) of reference sites (red bar), soil baselines are considered comparable (dark gray bar) and can be predicted based on changes in soil properties (black regression line). However, outside this area, the accuracy of predictions reduces (gray line). To determine the optimal radius for predicting soil health (here defined as pathogen density), pairwise changes were calculated using broad survey data (Fig. 1), generating a dataset comprising a total of 32,580 (181×180) sets of changes. **b: Changes in soil properties between random pairs of soils well explained plant health up to a distance of 70 km.** The pairwise changes dataset was filtered according to pairwise distance (steps of 10 km from 40 km to 3890 km) into the sub datasets and we measured the explanatory power of soil properties to plant health within each sub-dataset. The panel shows the decline in the explanatory power of the three soil properties, which best explain plant health as distance increases. For visualization, distances were divided into three classes: below 70 km (black), between 70 km and 330 km (dark grey), and more than 330 km (light grey). **c, Structural equation modelling showing different explanatory powers to plant health in different distance classes.** The SEMs with the highest explanatory power within each distance class are displayed. Black and red arrows indicate positive and negative effects, respectively. Numbers above each arrow and the arrow size represent the explanatory power of each path. Numbers below the box represent the total explanatory power of the models. The bar plot at the bottom represents the total explanatory power of the models. Abbreviations TN: soil total nitrogen, OM: soil organic matter content, PC1: coordinates of bacterial PCA analysis first axis, PD: pathogen density, Δ: changes (log-scale change proportion).

Next, we demonstrated that we could extrapolate our approach to predict plant health even without prior records of fertilization history. We hypothesized that geographically nearby sites would share similar agricultural management histories and would hence have comparable baselines for constructing plant health prediction models with prediction areas extending outward from these reference points (Fig. 3a). To validate our hypothesis, we computed pairwise changes in soil properties among all 181 soil samples from a broad sampling experiment (Extended Data Fig. 1a), generating a total of 32,580 sets of pairwise comparisons (Supplemental Table 11). Three key soil properties (nitrogen, organic matter defined as organic carbon and bacterial community composition) could explain plant health adequately (Extended Data Fig. 11). The explanatory power of these three soil parameters was best within a 70 km pairwise sample distance and decreased sharply beyond this threshold. Changes in soil total nitrogen content and organic matter explained up to 72% and 40%, of the plant health (Fig. 3b), respectively, indicating that nearby soils shared a comparable baseline for plant health. Notably, the explanatory power of bacterial PC1 showed the most evident distance-decay relationship, pointing to a strong dispersal limitation in soil^23^ (Fig. 3b and Extended Data Fig. 12). The SEMs integrating soil chemical and bacterial properties further showed that the explanatory power of models increased from 14.8% to up to 83.6% along with reduced pairwise sample distance within a 70 km radius (Fig. 3c and Supplementary Table 12). Based on these results, we used a stepwise model selection on samples within a 40 km distance to construct a spatial plant health prediction model, which predicted plant health with an accuracy of 83.6% when both soil chemical and microbiological properties were included (Extended Data Fig. 13). Collectively, our efforts resulted in models able to effectively predict plant health at local spatial scales, with prediction accuracy decreasing sharply along with increasing geographical distance.

Finally, we used our approach to predict temporal (Extended Data Fig.14), spatial (Fig. 4a), and spatiotemporal dynamics (Fig. 4b) in plant health based on high-resolution (0.1°×0.1° cells) historical soil property data (soil total nitrogen, organic matter, and pH) in China available in public databases from the years 1980 to 2015. Due to the lack of detailed fertilization records, we regard the period from 1980 to 2015 as representative of conventional farming practices, during which the use of chemical fertilizers in China increased substantially—from 12.7 million tons in 1980 to 60.0 million tons in 2014^24^. We first calculated the changes in soil chemical properties for each grid cell based on two constructed databases. We utilized the most parsimonious model (81.3% accuracy; see Material and Method) to predict temporal dynamics in plant health under conventional agricultural practice. In line with the widespread land degradation caused by intensive agriculture (FAOSTAT, Data of Access: [10-12-22]), we predicted a decline in plant health in most areas between 1980 and 2015 (average: -3.00%, median: -0.12% and 51.96% faced a decline, Extended Data Fig.14). We then estimated the annual change in plant health under organic farming scenarios based on the effects of soil properties from our meta-analysis (Fig. 2b). In contrast to conventional fertilization, we predict that transitioning back to organic fertilization could potentially improve plant health, reversing the annual decrease from -0.12% to a yearly increase of +1.78% (median value) and reducing the decline in plant health to only 30.78% of the estimated agricultural area (Extended Data Fig.14). We applied the same model to predict spatial dynamics in plant health using the broad survey experimental dataset, where known soil properties and plant health served as reference points (Extended Data Fig. 1a). Spatial predictions were calculated for each grid cell, focusing on areas within a 70 km radius of each reference site (with each site covering a 1° area). Using soil property data from 1980 and 2015, we modelled plant health around the reference sites and reproduced historical plant health scenarios of 1980 and 2015, respectively (Fig. 4a). Furthermore, we forecasted future plant health for 2050 under different fertilization scenarios combining this with the annual change in plant health (Fig. 4b). Under conventional management, plant health is projected to remain at low levels in over 60% of the areas included in the analysis up until 2050. In contrast, transitioning from conventional to organic fertilization could potentially enhance future plant health by 66%, with only 44.37% of the agricultural area predicted to experience low levels of plant health. The regions in northern China are estimated to benefit considerably in the future from the transition to organic farming, as indicated by both spatial and temporal predictions (Supplemental Figure 15).

**Fig. 4.**
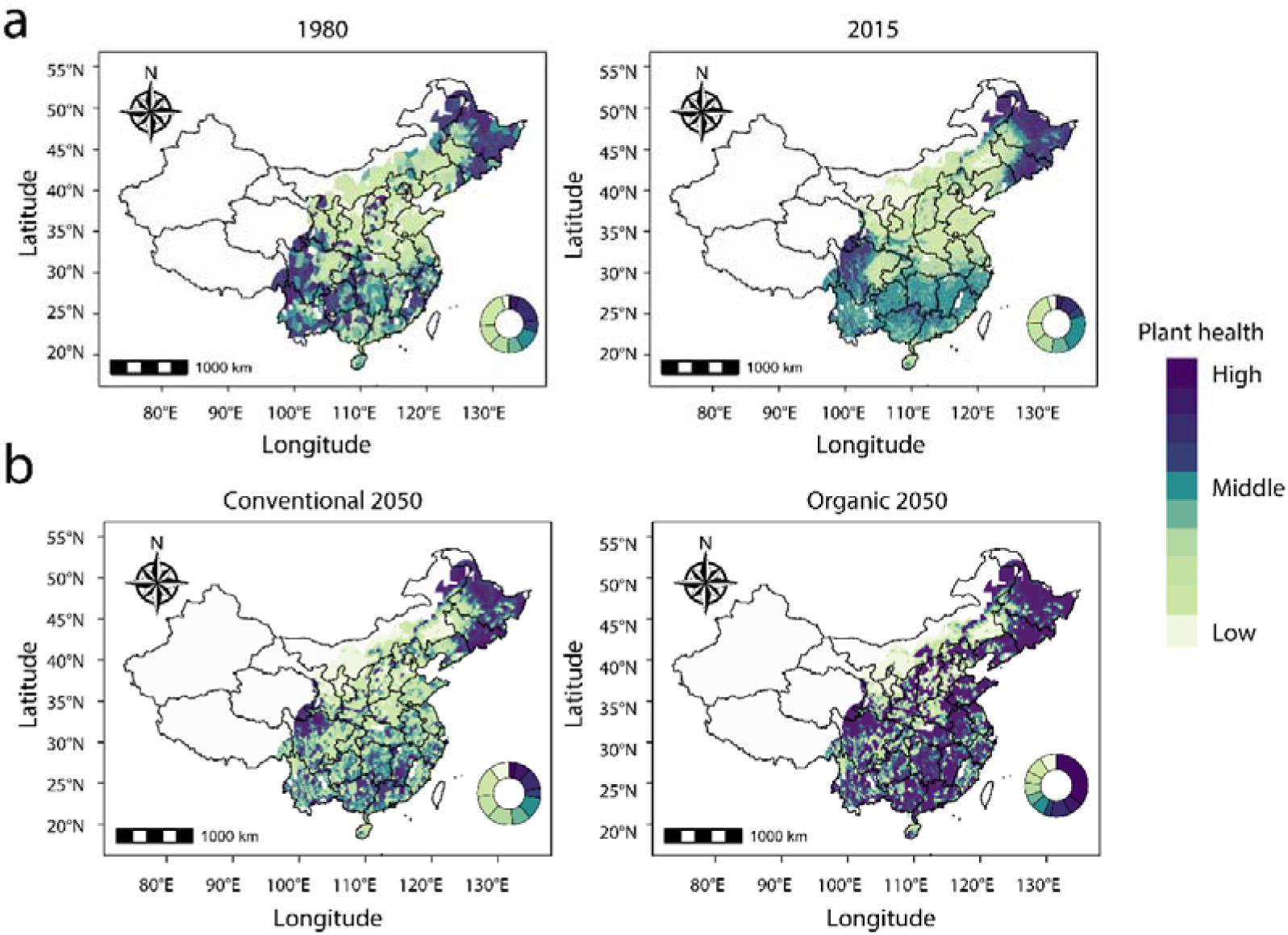
Spatiotemporal prediction of plant health in 1980, 2015 (a) and 2050 under conventional and organic fertilization scenarios (b). Cells with a resolution of 0.1° × 0.1° grids were generated to map soil health scenarios. The stepwise regression model with the best predictive accuracy (81.3%) was recruited for plant health prediction, with changes in soil chemical properties required as input. Soil properties data of cells were acquired from two public databases (Year 1980 and Year 2015) and calculated changes in soil properties between 1980 and 2015, which were used to predict soil health (here defined as pathogen density) changes under conventional scenarios. Improved soil properties induced by organic fertilization from meta-analysis (Fig. 2b) were extracted to estimate soil health under organic scenarios. The predicted changes in plant health were normalized by the time span (35 years from 1980 to 2015) and transformed into annual percentage changes for mapping. Leveraging 181 broad surveyed samples with both already known soil properties and plant health in the representation of the surrounding 1° radius area as reference areas, soil properties data from 2015 (the same year as the reference samples collected) were acquired for spatial prediction of plant health. According to the spatial extrapolation traits we discovered (Fig. 3b), plant health of cells within a 70 km distance around each reference area was predicted (Extended Data Fig. 13). Then we combined this spatial prediction with temporal prediction to estimate plant health in 2050 under conventional and organic fertilization scenarios (panel b). Predicted pathogen density was divided into ten classes for visualization, ranging from high (dark colors) to low (light colors). Circles on the map denote the proportions of each class for all predicted cells.

## Discussion

In this study, we present a new approach to predicting plant health based on changes in soil abiotic and biotic properties across varying spatial and temporal scales. Specifically, plant health responds consistently to organic farming-induced environmental changes despite having variable baselines that depend on geographic location and soil type. Using meta-analyses and empirical evidences, we demonstrate that the soil microbiota best predicts plant health and pathogen suppression and that soils low in organic matter and total nitrogen can greatly benefit from organic fertilization. In contrast, nutritionally rich soils are likely to benefit less from organic farming practices (Supplemental Figure 15), suggesting that more targeted approaches, such as the application of beneficial microbial inoculants^25^, might be needed to effectively improve plant health.

We found that no single set of soil properties could adequately predict plant health when challenged with pathogens across large scales, and none of the ten soil properties explained more than 5% of the variability in soil disease suppressiveness. However, plant and microbiome responses to organic farming were consistent across different soil types and climatic regions despite large changes in the baseline of soil suppressiveness between the sampling sites. This aligns well with previous studies showing that microbiome changes can predict the trajectory of plant health^26,27^. Specifically, we identified bacterial diversity as the most significant factor in explaining plant health, which has been previously associated with soil fertility and plant health by acting on nutrient cycling dynamics that affect nutrient availability in soils ^28^ and microbial competitive mechanisms that can enforce pathogen suppression through resource or niche competition^29^. The underlying benefits of microbial diversity are likely triggered by changes in soil chemical properties^30^, especially total nitrogen content. This could explain the improved explanatory power when both abiotic and biotic variables were included in the predictive models. For instance, the enhanced organic carbon in the soil can induce changes in the metabolic profile^31^ and functionality^32^ of microbial communities; these changes may create favorable conditions for more stable microbial communities^33^, enhancing their role in protecting plant health. These results lay the groundwork for predicting and manipulating plant health based on soil chemical and microbial properties using organic farming practices. However, such management practices would need to be tailored to each specific location and soil property baseline to achieve the desired changes and responses in plant health.

Our models could also be used to predict plant health across space. We could predict plant health up to a 70 km radius from the reference sampling sites based on microbial community composition and organic matter and nitrogen content. This is likely because nearby soils share similar agricultural management practices, climatic conditions and plant health baselines. However, we found that the predictive power of spatial models diminished with increasing geographic distance and that bacterial community composition was the most sensitive variable for spatial predictions. This could be explained by Tobler’s first law of geography and the biogeographical structuring of microbial populations in the soil that are inherently limited in their dispersal^34^. Also, other factors such as varying agricultural practices, climatic conditions, and soil characteristics could further reduce prediction accuracy over larger geographical ranges^35^. We also enabled temporary predictions of plant health by measuring changes in soil properties induced by differential management based on recorded history. Our findings suggest that the rate of change in soil properties, particularly microbial diversity driven by organic fertilization, is critical for understanding shifts in plant health. This highlights the importance of long-term management in shaping sustainable plant health outcomes. Additionally, our model indicates that future predictions of plant health could become more accurate over extended timeframes with continuous monitoring and the integration of real-time soil health data. Such data could guide the development of more adaptive soil management strategies tailored to the evolving needs of plants.

Here we highlight a few limitations of our models to guide further model development and experimental work. First, we measured soil suppressiveness only against the bacterial pathogen *R. solanacearum*. As such, extending this parameter to cover other and potentially multiple pathogens in agricultural systems ^36^ can likely improve the resolution of model predictions. Second, despite we found soil bacterial community properties to be important variables in the model, it is likely that the inclusion of other soil microbiota properties (e.g., parameters associated with the diversity and structure of fungal and protist communities) can provide a more comprehensive overview of the complexity of the soil microbiota, likely enhancing accuracy in the predictive models. Third, we focused our analysis on tomato plants as a model system. As such, caution is warranted in extrapolating these model predictions to other crop species, although fundamentals and parameters can be used as a reference to other datasets. Last, further information on the soil microbiota responses to multiple agroecosystem management strategies can also enhance complexity in model parametrization and data analysis. Overall, and despite these intrinsic limitations, our approach holds significant promise in predicting soil health and informing sustainable soil management strategies and restoration efforts. We demonstrate that plant health, a well-known complex concept in agriculture, can be modeled by using proper data parametrization and experimental systems. By distinguishing between baseline plant health and responses to environmental trends, our work offers a new perspective on soil health, emphasizing the need for dynamic management strategies. In conclusion, we provide a scalable and cost-effective method for monitoring and managing agricultural soil. This approach supports the development of high-yield, low-input sustainable farming systems.

## Methods

### Broad sampling experiment over 181 Chinese agricultural fields

A total of 181 randomized agricultural sites were selected across China, covering a broad range of soil types, climate regions and biomes (Supplementary Table 10). For all 181 sites, one soil sample from 0-20 cm was collected within the agricultural field. Soil chemical properties were measured following the Chinese standard for soil property determination (www.chinesestandard.net) using the following standards: pH (HJ 962–2018), soil organic matter content (NY/T 1121.6–2006) and total nitrogen content (LY/T 1228–2015). The catabolic potential of the soil microbiome was determined by integrating utilization capacity in 48 hours for 32 resources (Supplemental Table 1) following standard protocols^37^. Specifically, the resource use abilities were measured through principal component analysis, and the coordinates in the first component were extracted as the catabolic potential. The antifungal activity was defined as in vitro inhibition of soil microbial suspensions against *Fusarium oxysporum*. To determine the microbial properties of these soils, soil DNA was extracted using the MoBio PowerSoil™ DNA Isolation Kit (Mo Bio Laboratories Inc., Carlsbad, CA, USA) according to the manufacturer’s protocols from ∼0.5g soil samples. Subsequently, bacterial and fungal biomasses were defined using qPCR of 16S rRNA gene (primer Eub338/Eub518^38^) and ITS (primer ITS1F-ITS2R) copies, respectively, using SYBR Premix Ex Taq Kit (Takara, Dalian, China) according to the manufacturer’s instructions.

To determine bacterial composition in each soil sample, the V4-V5 region of the bacterial 16S ribosomal RNA gene was amplified by PCR (95 °C for 2 min, followed by 25 cycles at 95 °C for 30 s, 55 °C for 30 s, and 72 °C for 30 s and a final extension at 72 °C for 5 min) using primers 563F-802R with Illumina adaptors. Index sequences followed the standard PCR protocols. PCR reactions were performed in triplicate 20 μL mixture containing 4 μL of 5 × FastPfu Buffer, 2 μL of 2.5 mM dNTPs, 0.8 μL of each primer (5 μM), 0.4 μL of FastPfu Polymerase, and 10 ng of template DNA. Amplicons were extracted from 2% agarose gels and purified using the AxyPrep DNA Gel Extraction Kit (Axygen Biosciences, Union City, CA, U.S.) according to the manufacturer’s protocol. The sequence data were processed using the UPARSE pipeline^39^. Read pairs of each sample were assembled, and sequences were screened with the following criteria: 1) The 250bp reads were truncated at any site receiving an average quality score < 20 over a 10-bp sliding window, and the truncated reads shorter than 50bp were discarded. 2) reads containing ambiguous characters according to barcodes were removed. 3) only sequences that overlap over > 10 bp were assembled. After discarding the singletons, the sequence reads were clustered into operational taxonomic units (OTUs) at 97% similarity using the Deblur denoising algorithm. The 16S rRNA gene sequence was then analyzed by *uclust* algorithm (https://www.drive5.com/usearch/manual/uclust_algo.html) against the silva (Release 119) 16s rRNA database with 80% confidence threshold.

Plant growth and plant health were determined using greenhouse experiments. Nine-grid seeding pots were filled with 2 kilograms of soil that could contain nine tomato plants for each soil sample. Surface sterilized tomato seeds (Lycopersicon esculentum, cultivar “*Hongpen*”) were germinated on water agar plates and transferred to trays containing sterile seedling substrate. Tomato seedlings were incubated in the greenhouse at 30° ± 3°C for 3 weeks before transplanting to the pots. After three weeks, tomato seedlings were transplanted into nine-grid seeding trays, grown in a greenhouse (temperature range 25-35LJ) for 48 days, and watered with sterile water 3-7 times a week to maintain suitable growth conditions. Seedling pots with tomato plants were rearranged randomly every three days. Tomatoes planted in each soil were divided into two groups of trays for each soil, and each group contained nine tomato plants. The first group were cultivated until the fruiting stage to measure tomato plant biomass. Nine tomato plants without fruit were harvested and we measured the dry weight of harvested tomato plants. The second group was inoculated with *R. solanacearum QL-Rs1115* strain with a final density of 10^7^ CFU g/soil fourteen days after transplantation to measure plant health based on bacterial wilt severity and pathogen density. Twenty-five days after *R. solanacearum* inoculation, disease severity was recorded for all nine plants according to the percentage of wilted tomato plants for each soil sample^40^. Subsequently, rhizosphere soils from each cell were sampled, and DNA was extracted following established protocols. *R. solanacearum* numbers in the rhizosphere soil were quantified using qPCR with primer sets *Rsol_fliC*^41^. Pathogen Density (PD) was calculated by the ratio of *R. solanacearum* numbers to total bacterial gene copies.

### Meta-analysis of organic fertilization regime on soil properties and plant health

To investigate the effects of different fertilization regimes on soil chemical and microbial properties, we collected a total of 78 peer-reviewed papers published from April 2000 to April 2023 from Web of Science and Google Scholar to obtain a total of 135 independent experiments following PRISMA (Extended Data Fig. 3). These studies included one of the following two groups of keywords: “farmyard manure” “FYM” “manure(s)” “compost(s)” “composted” “straw” “organic waste(s)” “organic fertilizer(s)” “organic fertilization(s)” “organic amendment(s)” “biological amendment(s)” “organic farming” “organic garden” “organic agriculture” “organic input” “organic source” OR “microbial diversity” “microbial community” “bacterial diversity” “bacterial community” “fungal diversity” “fungal community” “soil biological” and “microbial properties”.

We selected representative datasets using the following criteria: 1) the study should include both organic and chemical fertilization regimes, with a similar or same amount of nitrogen input in the two treatments; 2) the study should include data on microbiome diversity (bacteria or fungi) and at least one of the following soil chemical properties: soil organic matter content, soil total nitrogen content and pH; 3) the duration of organic fertilization must have lasted ≥3 years; 4) depth of soil samples must be ≤20cm and 5) when the dataset contained multiple experiments, sub-datasets were considered as independent experiments if any of the site, soil type, crop type, organic fertilizer type, amount of nitrogen input, duration of fertilization, depth of soil samples or application methods differed. 6) Experiments with biochar as organic amendments were excluded. In addition to soil chemical and microbial properties, we collected basic information on the experimental sites, such as geographic coordinates (latitude and longitude), annual mean temperature, annual mean precipitation, crop type (at the time of sampling), organic content source, amount of nitrogen input and fertilization application duration. If annual mean precipitation and annual mean temperature were not included in the study metadata, they were extracted from the (http://www.worldclim.org/) database based on the study coordinates.

To investigate the effects of different fertilization regimes on plant health, we collected a total of 73 peer-reviewed papers from April 2000 to April 2023 from Web of Science and Google Scholar, and obtained a total of 202 independent studies following PRISMA (Extended Data Fig.3). These studies included one of the following two groups of keywords: “farmyard manure” “FYM” “manure(s)” “compost(s)” “composted” “straw” “organic waste(s)” “organic fertilizer(s)” “organic fertilization(s)” “organic amendment(s)” “biological amendment(s)” “organic farming” “organic garden” “organic agriculture” “organic input” “organic source” OR “disease(s)”, “pathogen(s)”, “suppression” and “immunity”.

We selected representative datasets fulfilling all the following criteria: 1) the study should include both organic and chemical fertilization regimes, with a similar or same amount of nitrogen input in the two treatments. 2) the study should include at least one type of data: diseased/healthy occurrence, disease incidence, disease index, disease severity or pathogen density. 3) when the dataset contained multiple experiments, sub-datasets were considered independent experiments if any of the sites, soil type, crop type, organic fertilizer type, amount of nitrogen input, duration of fertilization treatment, depth of soil samples or application methods differed. 4) Experiments with biochar as organic amendments were excluded. In addition, we also collected basic information on the experimental sites, such as geographic coordinates (latitude and longitude), pathogen species, host plant types, pathogen inoculation concentration, experimental site types and organic sources for further analysis and comparisons. When the datasets did not include geographic coordinates, those were estimated from Google Earth, according to the location of the experimental sites.

Experimental data were extracted from text, tables, or figures (using ‘g3data’ software). If the experiments reported only the standard error (SE), it was converted to the standard deviation (SD) using the following equation:

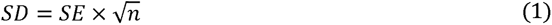

where n represented sample size. If the experiments showed no SE or SD, SE was set as 1/4 of the mean. For further analysis and comparison, measurements of soil organic carbon (SOC, g·kg^-1^) were considered equivalent to soil organic matter (OM, g·kg^-1^) using the following equation:

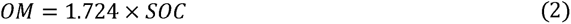

pH-CaCl_2_ was transformed to pH-H_2_O using the following equation:

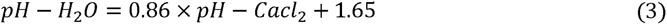

The database containing observations of survival or dead plants record were transformed into disease severity (*DS*) using the following equation:

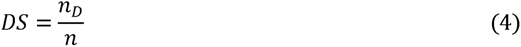

or

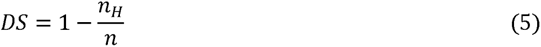

where *n_D_* represented diseased sample size, *n_H_* represented healthy sample size and *n* represented total sample size. The standard error of these transformed data was set as 1/4 of the mean.

The ratio of soil properties and plant health (disease severity and pathogen density) was calculated using the following equation:

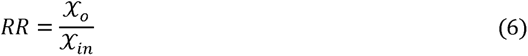

where *RR* represent the response ratio of soil properties and plant health, *X*_O_ and *X_in_* represented soil properties and plant health in organic and inorganic (conventional) regime soil, respectively.

For comparison, the variance of response ratio (*v*) was calculated using the following equation:

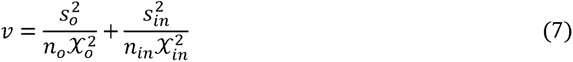

Where *s_O_* and *s_ln_* represent standard deviations of soil properties and plant health in organic and inorganic soil, respectively. *n*_O_ represents sample size in organic soil, *n_ln_* represents the sample size of inorganic soil.

The response ratio was weighted using the following equation:

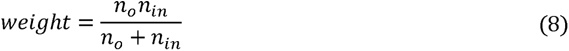

Changes (*Δ*) of soil properties and plant health was calculated using the following equation:

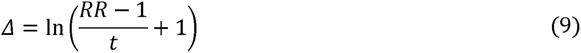

where *t* represented the time span (duration) of the organic fertilizer regime. OpenMEE^42^ was used to calculate the weighted response ratio.

### Paired-sampling experiments in 23 Chinese agricultural fields

From May to July 2018, soil samples were collected from 23 agricultural fields across China covering various climate and soil types. Each site contained three replicates of organic and nearby inorganic fertilization regime samples, respectively. All organic fertilization treatments lasted for at least five years. A total of 138 soil samples were obtained with their geographic coordinates, and the duration of organic fertilization was recorded. Soil properties were measured following the same methods as in the broad sampling experiment described earlier.

Soil microbial properties were determined by metagenomic sequencing. Microbial DNA was also extracted following a method consistent with the one used in the broad sampling experiment. Total soil DNA (∼450bp) was ligated to Illumina adaptors using the TruSeq™ DNA Sample Prep Kit and shotgun-sequenced in Illumina Hiseq platform at Shanghai Biozeron Biological Technology Co. Ltd (China). After building the PE library by Illumina HiSeq 4000, 150-bp paired-end technology, raw data was quality checked Trimmomatic^43^ and removal of host DNA was conducted by Bowtie2^44^, the rationality and effectiveness of quality checking were estimated by FastQC^45^. Reads were assembled using Megahit^46^ into contigs and predicted genes by METAProdigal^47^. A non-redundant gene catalogue was built based on predicted gene sequence by CD-hit with 95% identity and 90% coverage. Based on the non-redundant gene catalogue, microbial composition annotation was performed by NCBI-nr database. Plant health was measured following a method consistent with the one used in the broad sampling experiment.

### Calculation of pairwise changes between paired soil samples

Changes in soil properties and plant health between all 181 pairwise samples were calculated, resulting in a total of 32,580 (181 samples ×180 pairs) pairwise comparisons. As time span was the random factor, changes were calculated according to equation (9), and *t* was set as 1. A linear fitted model was then used to measure associations between soil properties and pathogen density, which were used as a proxy for plant health. To investigate the impact of distance on the explanatory power, we filtered the changes dataset according to pairwise geographical distance, ranging from 40 km to 3,890 km with 10 km steps, thereby generating distinct sub-datasets corresponding to distance thresholds. Adjusted R-square values from the linear regression models were utilized for each sub-dataset to mitigate the influence of degrees of freedom when evaluating explanatory power. Geographical distances were categorized into three classes based on distance-explanatory power curves: close (less than 70 km), intermediate (70 km to 330 km), and distant (more than 330 km). Finally, the structural equation and stepwise regression models followed the previously mentioned steps and were constructed to evaluate predicted accuracy within each distance class sub-dataset.

### Modelling spatiotemporal prediction of plant health

The step-wise model constructed in each sub-dataset of the corresponding pairwise distance thresholds enabled us to assess a spatial range of the model for estimating spatial scenarios of plant health. A spatial range of 70 km for the model was selected and optimized to attain broad spatial coverage while preserving precision to the greatest extent possible. The model utilizes changes in soil properties (pH, organic matter content and total nitrogen content) relative to reference sites as inputs and can predict changes in pathogen density. Subsequently, we estimated spatial scenarios of plant health on a 0.1×0.1-degree resolution with the following existing regional data sets: (1) Basic soil property dataset of high-resolution China Soil Information Grids^48^ (2015, http://www.geodata.cn) from National Earth System Science Data Center, National Science & Technology Infrastructure of China (http://www.geodata.cn). (2) Native database in China based on the second national soil survey data sets^49^ (1980s, http://www.sciencedb.cn/dataSet/handle/88).

To predict temporal dynamics in plant health, we first extracted soil properties data from public dataset in 1980 and 2015, then calculated Changes (*Δ*) of soil properties between 1980 and 2015 using Eq.9, these changes in soil properties were input to the model to calculated Changes of pathogen density. Changes (*Δ*) of pathogen density were transformed into annual percentage change of pathogen density (*r*) using the following equation:

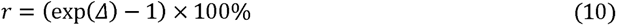

To estimate the annual change in plant health under the organic fertilizer regime, the weighted changes in soil properties from the meta-analysis were used as input in the stepwise regression model to predict changes in pathogen density induced by organic fertilization. The estimate annual change of plant health under organic fertilization (*r_o_*) the following equation was used:

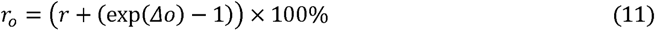

Where *Δo* represented output changes in pathogen density induced by the organic fertilizer regime from the stepwise model.

To predict spatial dynamics of plant health in the year 2015, we extracted soil properties data from the public dataset in 2015 and regarded our broad sampling experiment as reference sites to calculate changes in soil properties as inputs of the model. The included cell followed these criteria: 1) the cell will be calculated soil properties and plant health changes relative to the nearest reference site. If no reference site were found within a 70 km radius, the grid was excluded from the map. 2) soil properties data from 5∼15cm depth were used, which is same sampling depth used in broad sampling experiment. The spatial dynamics of plant health in year 1980 were calculated according to annual changes of plant health based on spatial dynamics of plant health in year 2015.

To map spatial scenarios of plant health in year 2050 under conventional fertilization regime, we used assumption that the pathogen effect on plant health between 1980 to 2015 was consistent, and thus, annual percentage change of pathogen density followed Eq.10; the spatial scenarios of plant health in year 2050 under organic fertilizer regime were estimated using estimated annual percentage change of pathogen density under organic fertilizer regime generated from Eq.11.

For all spatial scenarios of plant health, the pathogen density was normalized by the baseline of each grid to generate plant health levels. Baseline of pathogen density (*β*_0_) was mathematically defined as the value when total nitrogen and soil pH equal one, which we could represent using the following equation (Extended Data Fig. 1):

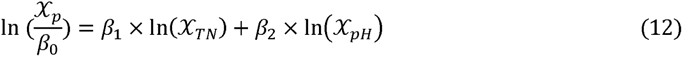

Where x_p_ represent pathogen density pathogen density, x_TN_ represent total nitrogen content and x_pH_ represent soil pH of each grid, and *β*_1_ represent the coefficient of total nitrogen changes￼ represent the coefficient of pH changes in the stepwise regression model. This equation can be transformed to calculate baseline of pathogen density as:

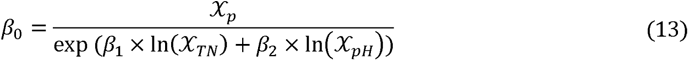

Estimated spatial scenarios of plant health in the year 2015 were used to calculate the baseline.

### Statistical analysis

All statistical analyses were performed in R version 4.3.2. Function ‘*lm*’ was used to build a linear fitted model between soil properties and plant health, and the extracted r-square value was used as the measure of explanatory power. Leverage analysis was conducted for outliers to exclude the leverage effect of a single data point by package ‘car’ (Supplementary Table 6). To reduce the classification error of microbial taxonomy based on non-redundant gene catalogue annotation, genus level was selected for further analysis. Bacterial and fungal diversity (Shannon index, richness) were calculated using the *vegan* package^50^. Bacterial and fungal community structure was assessed through PCA analysis and the first principal component coordinates were extracted using the *prcomp* function. Changes in soil properties and plant health were calculated following equation (6)-(11). Linear fit modelling by function ‘*lm*’ was used to evaluate associations between soil properties and plant health, and then r-squared was extracted as the measure of explanatory power. Genera with changes in relative abundances were significantly associated with both disease index and pathogen density and were identified as disease-correlated bacterial genera.

The structural equation modelling (SEM) was constructed using the package ‘lavaan’^51^ to evaluate the effects of soil chemical and microbial properties and their interactions with plant health. All data used in the SEM was first normalized by function ‘scale’ before the construction.

Statistics of SEM were extracted by ‘semTools’ packages and the overall goodness of fit was assessed, and the model was further considered when the following criteria were met: ratio of χ2 and degree of freedom <2, RMSEA (root mean square error of approximation) ≈ 0 and GFI (goodness-of-fit index) >0.80. A stepwise regression model was used to identify the most parsimonious set of predictors based on the AIC of the model. All significantly associated factors were first included in the model, and then we used a stepwise algorithm in both directions searching mode using the function ‘step’. Specifically, for changes, the intercept was remove when constructing a stepwise regression model. After the best model was constructed, the equation of the model was extracted for prediction and measurement of prediction accuracy.

## Methods

The meta-analysis database can be accessed in supplementary tables 1 and 2. The metagenomic raw sequences data were uploaded to NCBI under BioProject ID PRJNA912236, BioSample accession SAMN32235789 to SAMN32235926. The 16S rRNA amplicon raw sequences were uploaded to NCBI under BioProject ID PRJNA933245, BioSample accession SAMN33230199 to SAMN33230379.

## Acknowledgements

We thank G.R. Shen for the supporting the greenhouse experiment and K.M. Yang, Y.Z. Zhang, and X.R. Yang for technical assistance. This research was supported by the National Key Research and Development Program of China (2021YFD1900100), the National Natural Science Foundation of China (41922053, and 31972504).

## Contributions

Z.W. conceived the project and designed the experiments. Z.W., Y.D.M., C.Q.L., P.J.C and N.Q.W. collected the soil samples. Z.W., Y.G.H and N.Q.W. conducted the experiment. N.Q.W. performed the statistical analysis with the help of A.J., Z.W. and G.F.J. Z.W., N.Q.W., A.J. and G.F.J. wrote the first draft, all authors discussed and commented on the results and on the manuscript.

## Ethics declarations

The authors declare no competing interests.

