## Extended Data Fig.1-15 for "Spatiotemporal changes in soil properties predict plant health at continental scale"

### Title

^1^ Jiangsu Provincial Key Lab for Organic Solid Waste Utilization, National Engineering Research Center for Organic-based Fertilizers, Jiangsu Collaborative Innovation Center for Solid Organic Waste Resource Utilization, Nanjing Agricultural University, Weigang 1, Nanjing, 210095, PR China. Laboratory of Soil Health, Nanjing Agricultural University, Nanjing 210095, China

^2^ Nanjing Institute of Environmental Sciences, MEE, 8 Jiang-Wang-Miao Street, Nanjing, 210042, China

^3^ National Agricultural Technology Extension and Service Center, Beijing, 100125, China

^4^ Setec Energie Environnement. 97/101 Bvd Vivier Merle. 69003 Lyon. France

^5^ Department of Microbiological Sciences, North Dakota State University, Fargo, North Dakota, USA

^6^ Freie Universität Berlin, Institute of Biology, D-14195 Berlin, Germany

^7^ Berlin-Brandenburg Institute of Advanced Biodiversity Research (BBIB), D-14195 Berlin, Germany

^8^ Department of Plant Science & Huck Institutes of the Life Sciences, The Pennsylvania State University, University Park, PA, USA

^9^ The One Health Microbiome Center, Huck Institutes of the Life Sciences, The Pennsylvania State University, University Park, PA, USA

^10^Department of Microbiology, University of Helsinki, 00014, Helsinki, Finland

### ^*^Correspondence

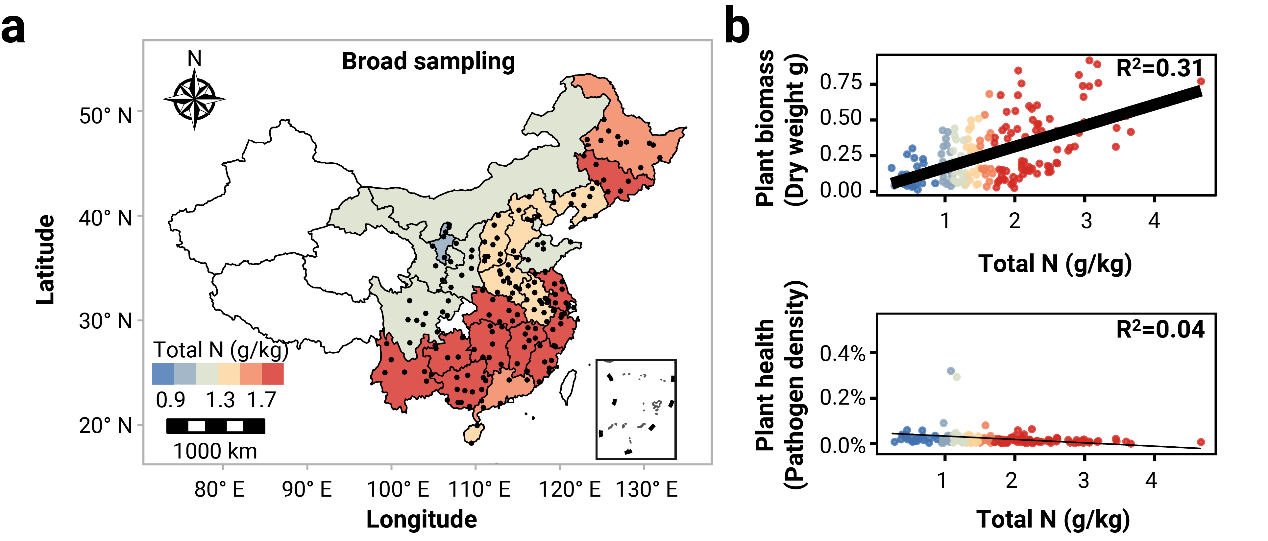

**Extended Data Fig.1 Broad sampling evaluating difficulties in predicting plant growth and plant health. a: Geographical overview of the 181 soil samples collected in China.** Each point corresponds to a collected soil sample. For each point, we measured a range of soil properties (Table 1) and ecosystem functions associated with plant growth (tomato dry weight), soil biological activity (soil bacterial and fungal biomass, catabolic potential, antifungal activity), especially the ability to maintain plant health under pathogen pressure (pathogen *Ralstonia solanacearum*, pathogen density and disease severity). The fill colors denote the average total nitrogen content (g/kg) across sampling sites within province. **b: Soil total nitrogen predicts well tomato biomass in absence of disease, but do not predict plant health, in term of pathogen *Ralstonia solanacearum* density in tomato rhizosphere*.*** As a best explanatory factor among all variables (see all in Supplemental Figure 1), nitrogen was selected as an example. Colors denote the measured total nitrogen content of soils.

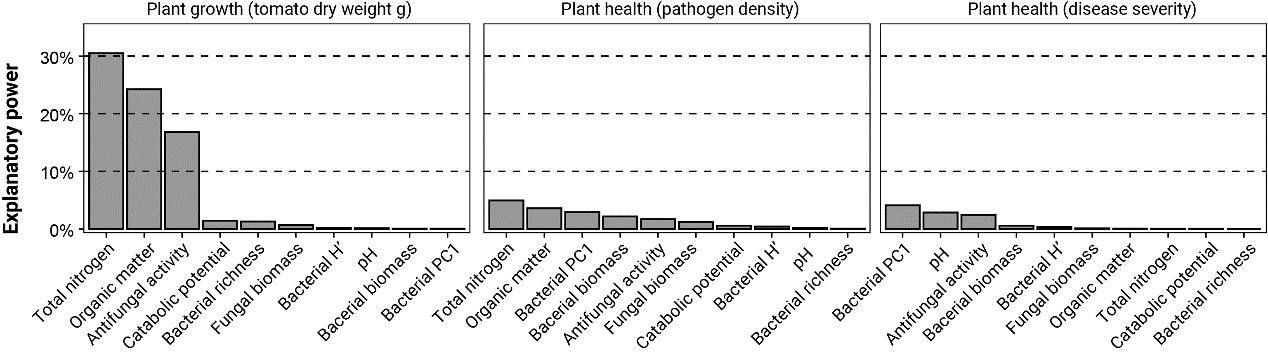

**Extended Data Fig.2: Snapshot measurements of soil properties only capture plant growth (plant biomass), but not plant health.** Linear fitted models were recureited to assess expalnation poweres of soil properteis to plant growth and health. Plant growth was defined as dry weight of tomato plant (in flowering stage without fruits) in absence of disease.Plant health was defined as soil suppresivenss (pathogen density and disease severity). Pathogen density was measured as the proportion of qPCR on fliC (pathogen qPCR) to total bacterial biomass, Disease severity was defined as the average disease index of tomato plants after soil inoculation with *R. solanacearum*. For soil properties, antifungal activity was defined as in vitro inhibition of soil suspension against Fusarium oxysporum; Catabolic potential was defined as integrated utilization capacity for 32 carbon sources; Bacterial and fungal biomass were defined by qPCR on 16S rRNA gene and ITS, respectively; Bacterial richness, H’ (Shannon index) and PC1 (the first princple compotement of community ) were defined by 16s rRNA amplicons.

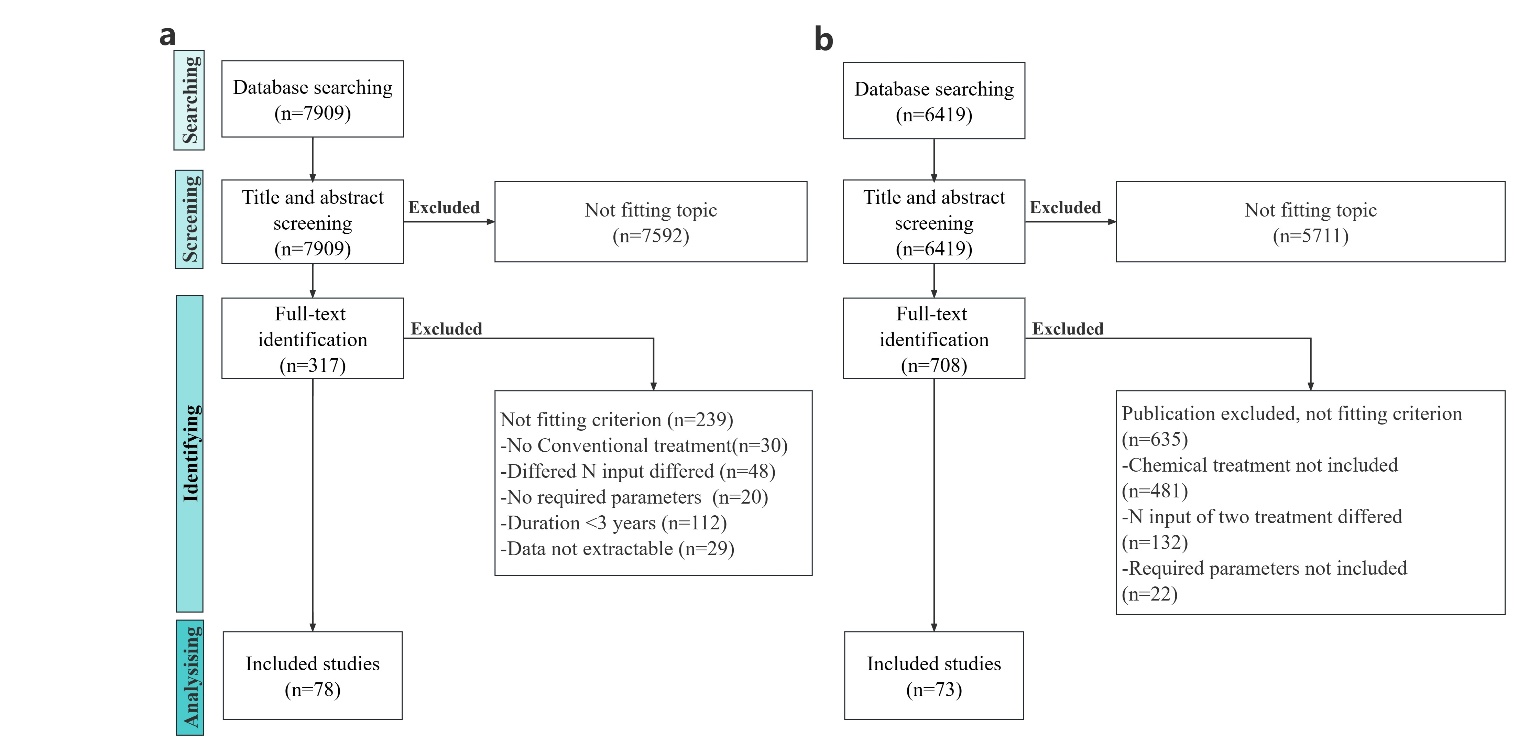

**Extended Data Fig.3:** **The PRISMA diagram of meta-analysis workflow.** **a, Meta-analysis to investigate the effects of different fertilization regimes on soil chemical and microbial properties. b, Meta-analysis to investigate the effects of different fertilization regimes on soil immunity.** Numbers in brackets represent studies numbers included in each step. The studies were excluded following the meta-analysis criteria (see Methods for details).

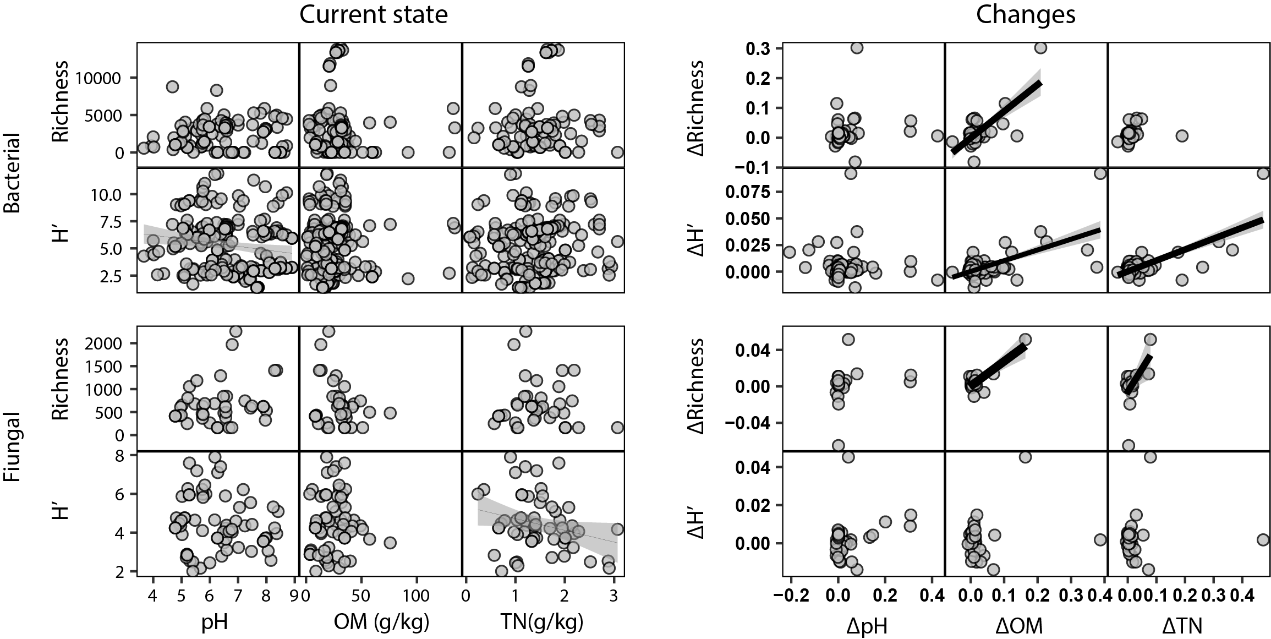

**Extended Data Fig.4: Associations between soil chemical properties and microbial properties in terms of current state (left) and changes (right) in meta-analysis.** Each point corresponds to one tested soil samples (for current state) or changes between one pair of soil samples (for changes). Line size represents explanatory power of chemical properties to microbial properties. The shaded area represents the 95% confidence intervals (CI).

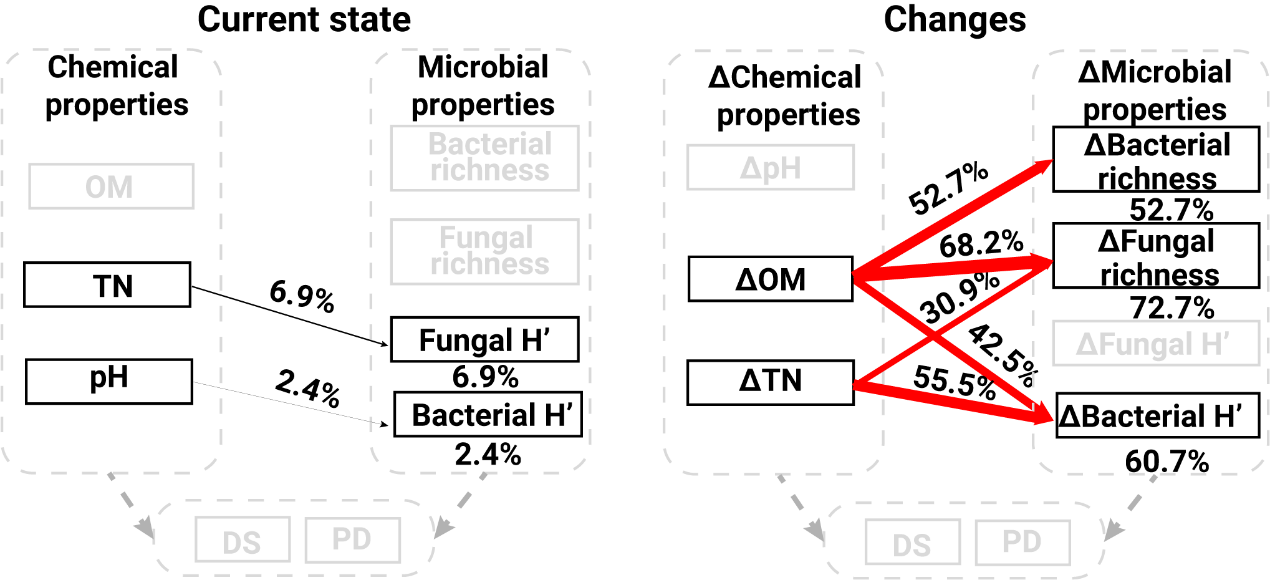

**Extended Data Fig.5: Graphical overview displaying the explanatory power of soil chemical properties on microbial properties (bacterial and fungal) expressed as a function of current soil characteristics (left) and changes (right).** Changes was defined as the log-transformed ratio of the assessed variable normalized by time span between the organic and chemical fertilizer treatments in each paired observation. Black and red arrows indicate positive and negative effects, respectively. Percentage values displayed above each arrow and arrow size represent the explanatory power of the respective paths. The numbers below each response variable summarize the total explanatory power of all significantly associated variables. Abbreviations, TN: soil total nitrogen content, OM: soil organic matter content, H’: Shannon index, DS: disease severity, PD: pathogen density, Δ: changes.

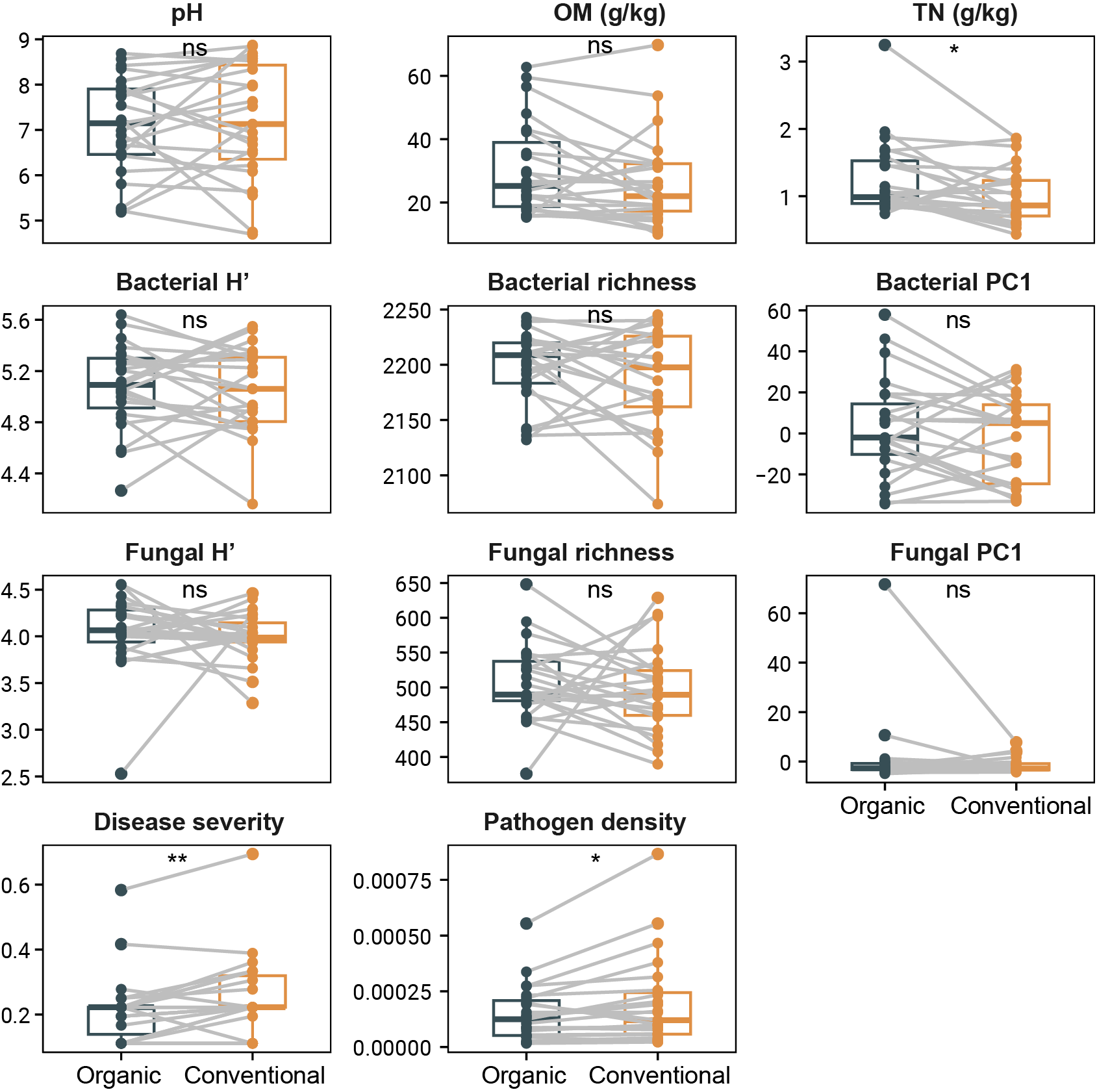

**Extended Data Fig.6: Boxplot illustrating pairwise differences in soil properties and plant health (disease severity and pathogen density) between organic and conventional fertilization in pairwise sampling.** Dots represent average values of corresponding soil properties or plant health in each sampling site. Asterisks represent p value of paired t test. ns: p>0.05, *: 0.05>p>0.01, **: 0.01>p>0.001.

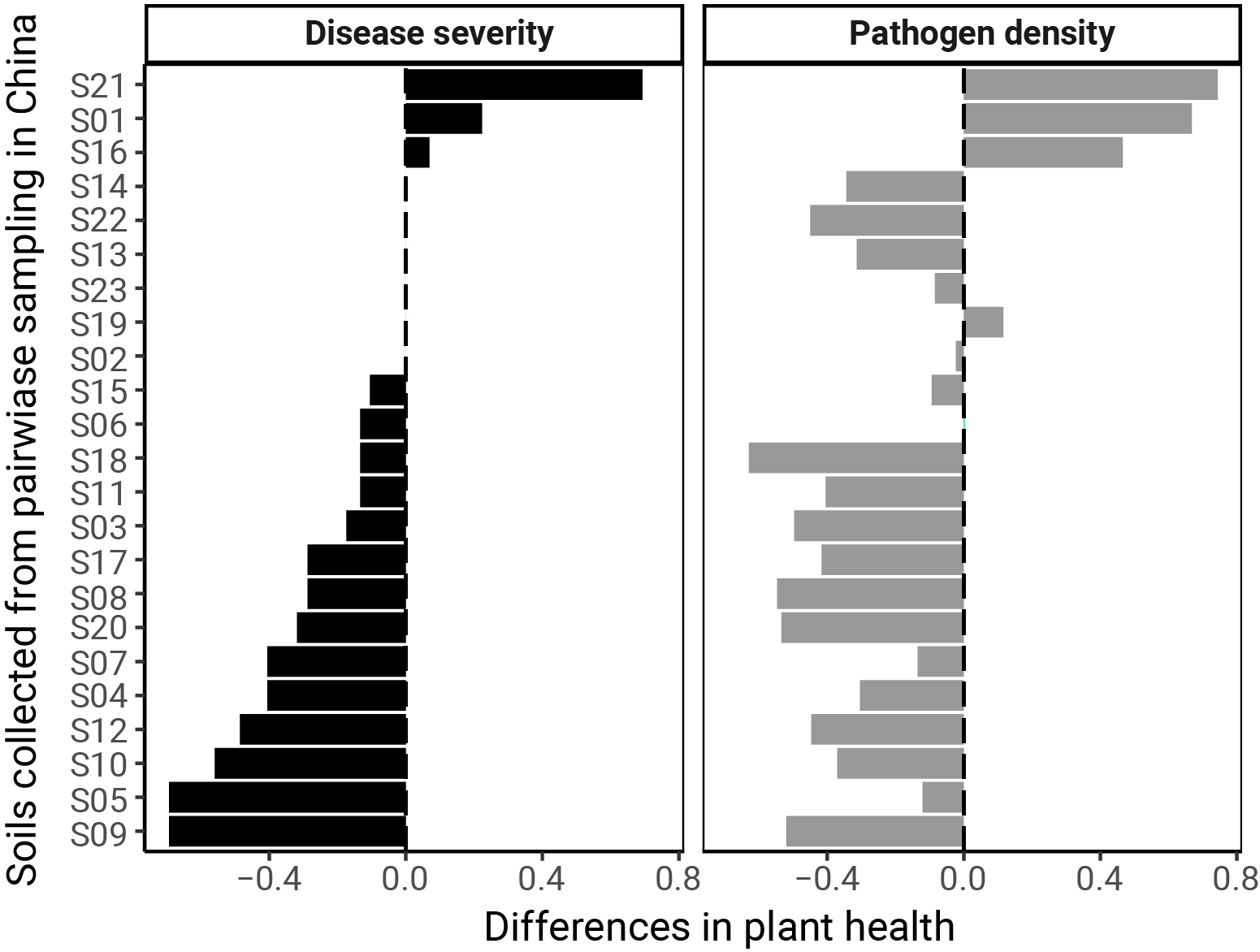

**Extended Data Fig. 7: Differences in plant health (disease severity (DS) and pathogen density (PD)) led by fertilization regime (Organic vs inorganic) in 23 pairwise sampling agricultural fields.** Differences was calculated by log-proportion values of plant health between organic and inorganic fertilization. X axis represents differences in plant health, which was defined as the log-transformed ratio of pathogen density and disease severity between two paired samples at one site (organic / inorganic), Y axis represents sampling site ID. Differences above 0 represents organic fertilizer regime increased disease severity or pathogen density.
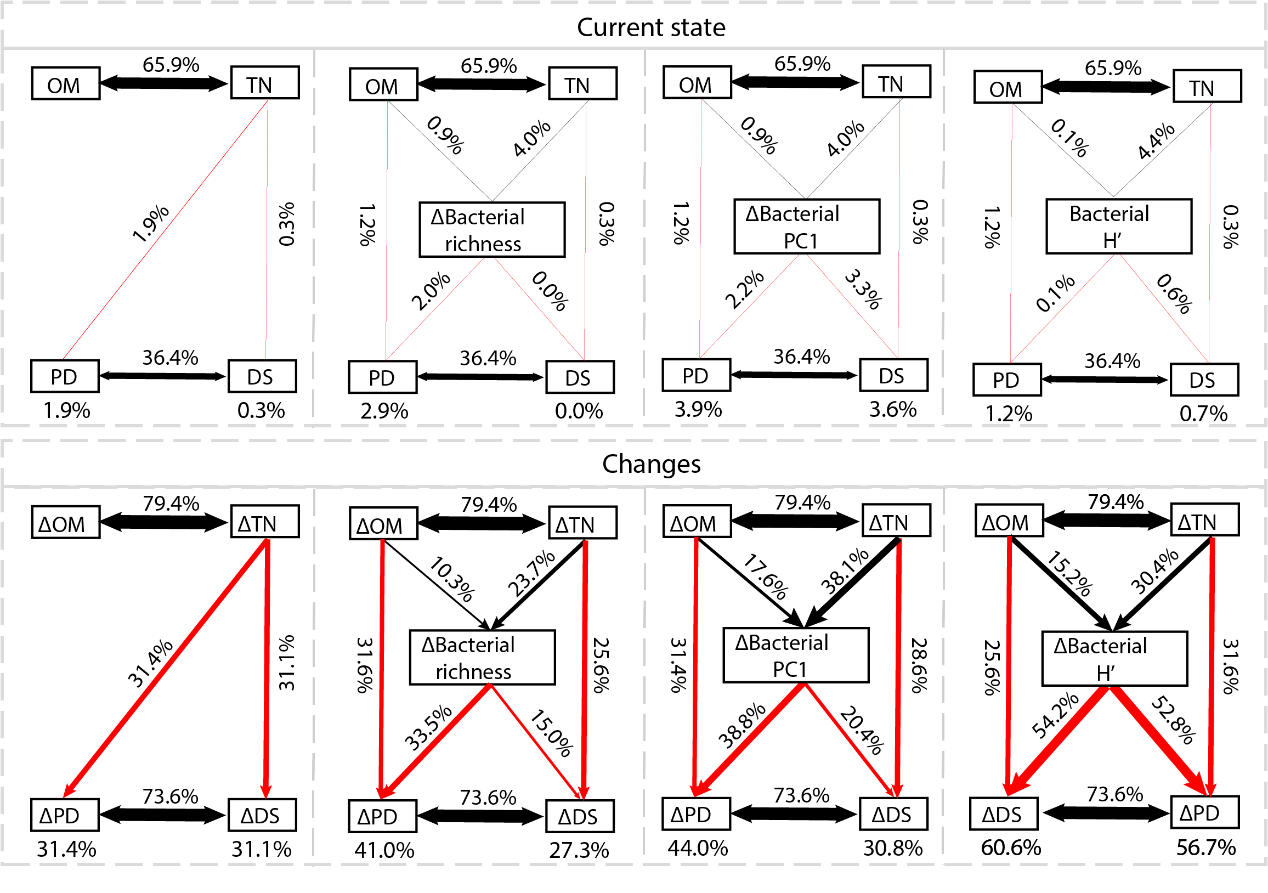

**Extended Data Fig. 8: Structural equation modeling highlighting the key relationships between chemical properties, microbiome composition, and plant health.** The above panel shows the effect of snapshot measurement (current state), the bottom panel shows the effect of changes. Black and red arrows indicate positive and negative correlations, respectively. Percentages above each arrow and the arrow size represent the explanatory power of each path. The number at the bottom represents the total explanatory power of the SEM to plant health. Abbreviations, TN: soil total nitrogen content, OM: soil organic matter content, H’: Shannon index, PC1: first axis of the PCA analysis on bacterial community, DS: disease severity, PD: pathogen density, Δ: changes.

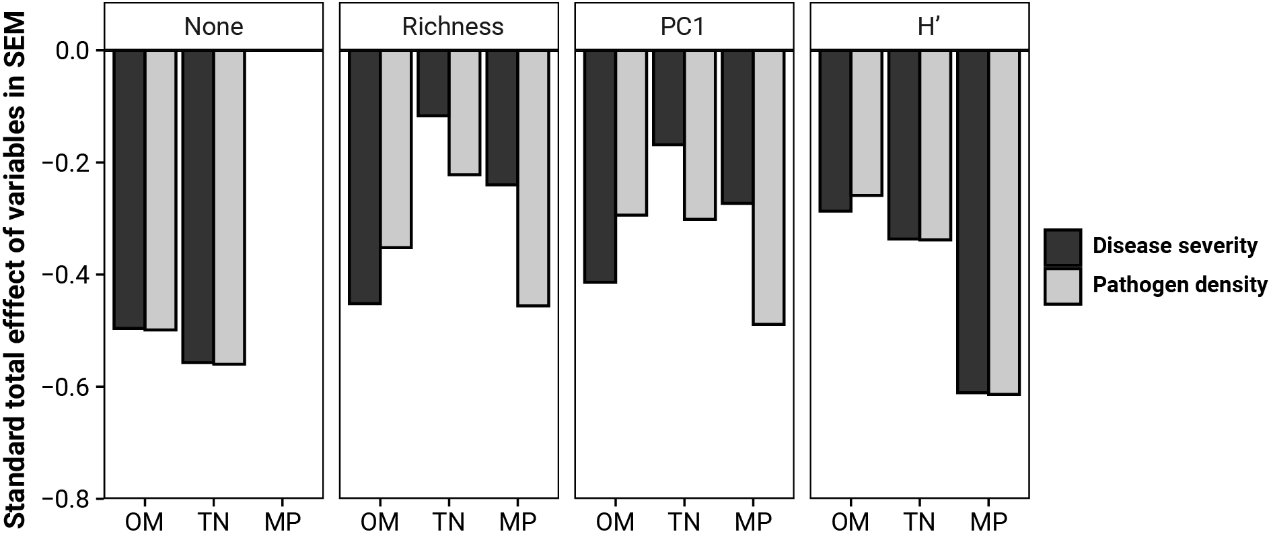

**Extended Data Fig. 9:** **Standard total effect of variables to plant health (disease severity and PD: pathogen density) in structural equation modeling (SEM) in term of changes.** Labels in facet represents in included microbial variables in SEM. None: no microbial variables included. Richness: bacterial richness, PC1: bacterial PC1, H’: Bacterial Shannon index. Abbreviations, OM: soil organic matter content, TN: soil total nitrogen content, MP: microbial properties

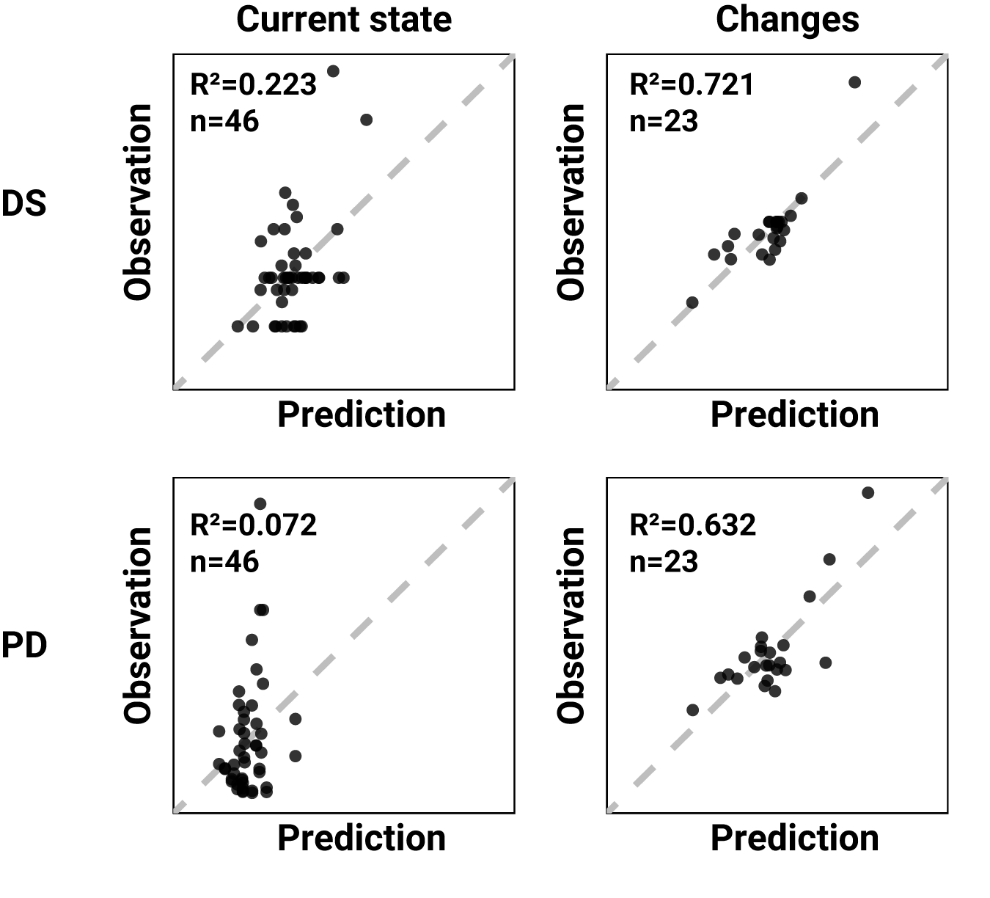

**Extended Data Fig. 10: Predictive accuracy of plant health (disease severity and pathogen density) according to soil properties based on current state (left) and pairwise sample changes (right) of soil parameters.** The most parsimonious set of parameters was selected using a step-wise model selection with bidirectional data dropout starting from an empty. X-axis represents prediction value predicted by step-wise regression model; y-axis represents observation value of plant health from experimental measurements. Gray dotted line represents a standard curve of slope=1.

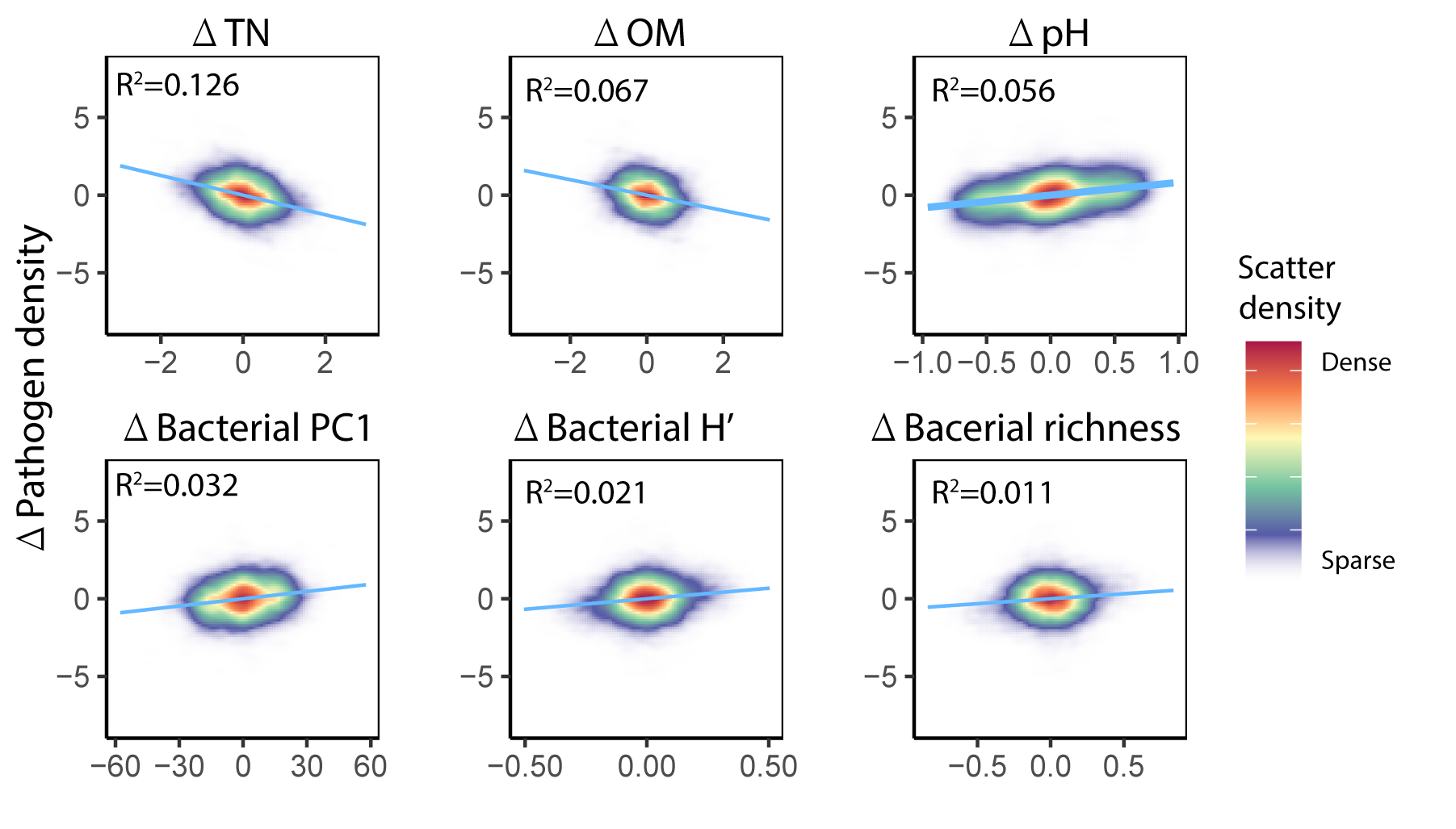

**Extended Data Fig. 11: Scatter density plot showing associations between each soil properties and pathogen density in terms of changes.** Pairwise changes in soil properties and pathogen density among all 181 soil sampling sites were calculated, which generated a total of 32580 sets of pairwise changes dataset. Due to the large scale of dataset, scatter plot was transformed into scatter density plot for better visualization. Colors of area represent the scatter density. X-axis represents changes of each soil characteristics; Y-axis represents changes in pathogen density.

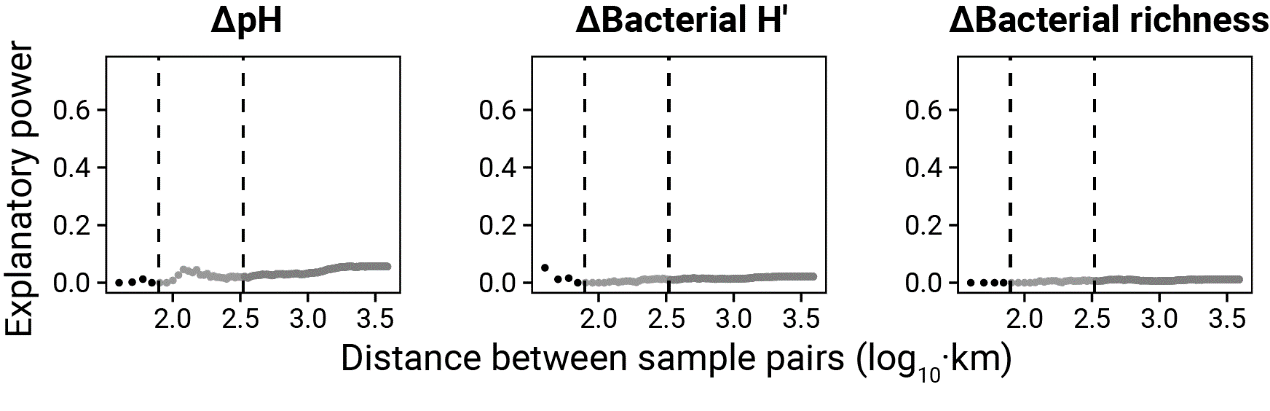

**Extended Data Fig. 12: Explanatory power of three soil properties (pH. Bacterial H’ and richness) to plant health decline with the distance between sample pairs increase.** The pairwise changes dataset were filtered according to pairwise distance (steps of 10 km from 40 km to 3890 km) into sub dataset and measured explanatory power of soil properties to plant health within each sub dataset. The panel shows the decline in explanatory power of the other three soil properties to plant health as distance increases. For visualization, distances were divided into three classes: below 70 km (black), between 70 km and 330 km (dark grey), and more than 330 km (light grey).

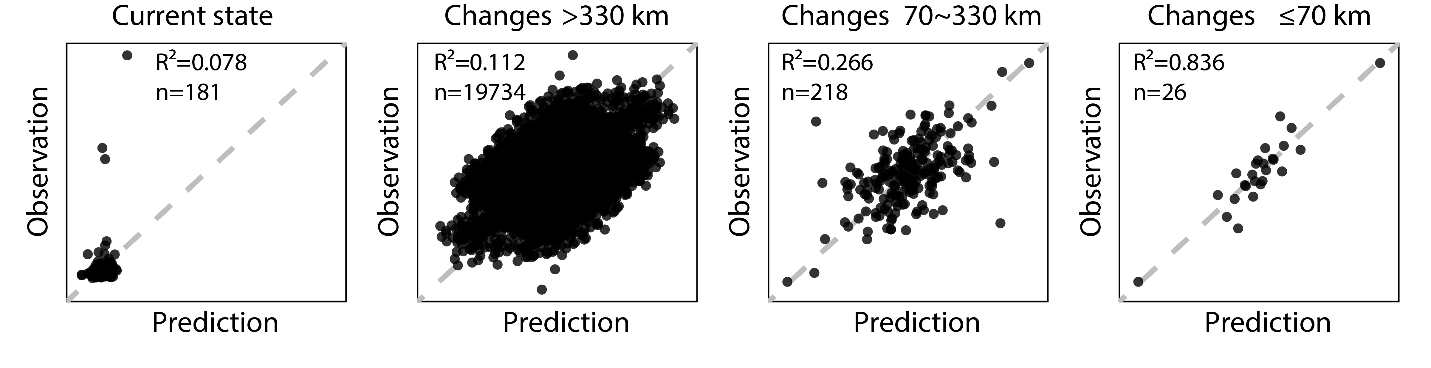

**Extended Data Fig.13: Predictive accuracy of plant health as a function of soil parameters used as current state and their changes at different pairwise distance threshold.** The most parsimonious set of parameters was selected using a step-wise model selection with bidirectional data dropout starting from an empty. X-axis represents prediction value predicted by step-wise regression model; y-axis represents observation value of plant health from experimental measurements. Points represent individual samples (current state) or pairs of soils (changes). Gray dotted line represents a standard curve of slope=1.

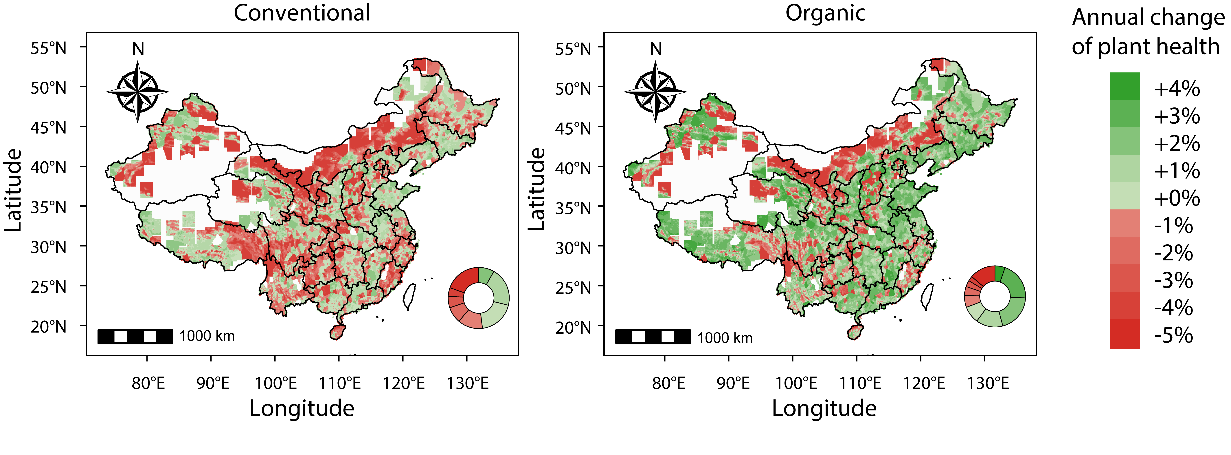

**Extended Data Fig.14: Temporal prediction of plant health under conventional and organic fertilization scenarios.** Cells with a resolution of 0.1° × 0.1° grids were generated to map soil health scenarios. The stepwise regression model with best predictive accuracy (81.3%) were recruited for plant health prediction, with changes in soil chemical properties were required as input. Soil properties data of cells were acquired from two public database (Year 1980 and Year 2015) and calculated changes in soil properties between 1980 and 2015, which were used to predict soil health (here defined as pathogen density) changes under conventional scenarios. Improved soil properties induced by organic fertilization from meta-analysis (Fig. 2b) were extracted to estimate soil health under organic scenarios. The predicted changes in plant health were normalized by the time span (35 year from 1980 to 2015) and transformed into annual percentage changes for mapping. Circles on the map denote the proportions of annual percentage change of plant health for all predicted cells.

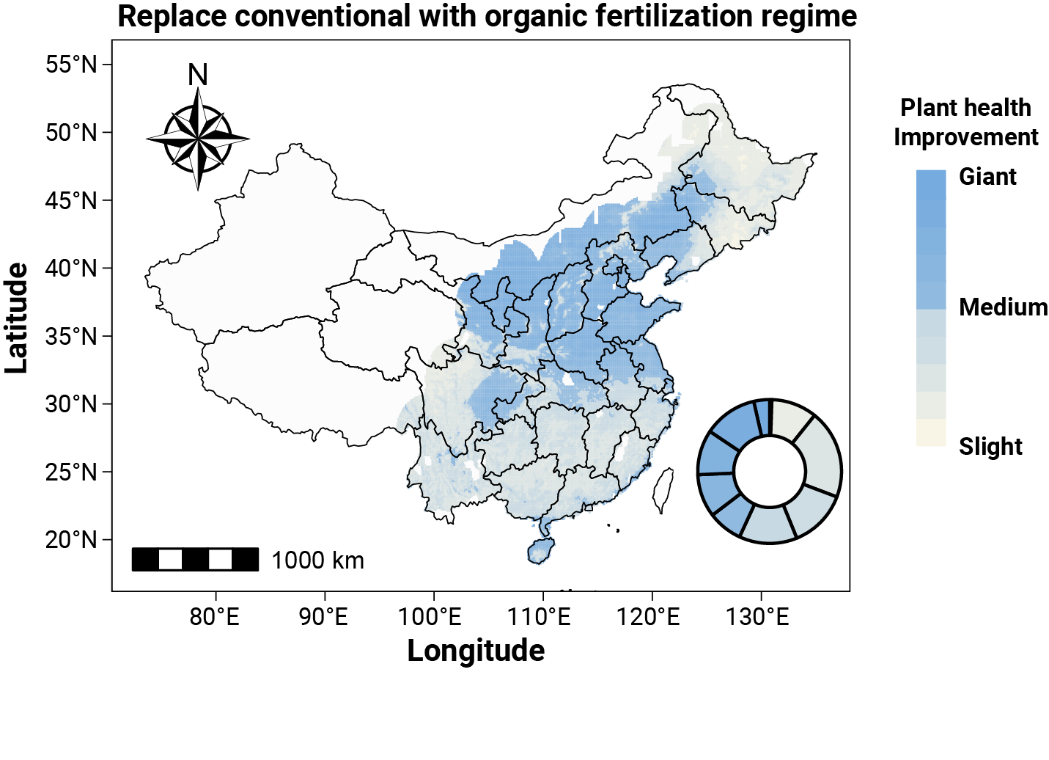

**Extended Data Fig.15:** **Estimated improvement in plant health replaced conventional with organic fertilization regime to 2050.** Improvements in plant health were assessed by differences in absolute value of pathogen density between organic and conventional fertilizer regime estimated in 2050. Improvements were divided into ten classes for visualization. Colors represent ten plant health improvement classes from high (dark) to low (light). Circles on the map represent proportions of each class for all predicted cells.
